# ESCA pipeline: Easy-to-use SARS-CoV-2 genome Assembler

**DOI:** 10.1101/2021.05.21.445156

**Authors:** Martina Rueca, Emanuela Giombini, Francesco Messina, Barbara Bartolini, Antonino Di Caro, Maria R. Capobianchi, Cesare E. M. Gruber

## Abstract

Early sequencing and quick analysis of SARS-CoV-2 genome are contributing to understand the dynamics of COVID19 epidemics and to countermeasures design at global level. Amplicon-based NGS methods are widely used to sequence the SARS-CoV-2 genome and to identify novel variants that are emerging in rapid succession, harboring multiple deletions and amino acid changing mutations. To facilitate the analysis of NGS sequencing data obtained from amplicon-based sequencing methods, here we propose an easy-to-use SARS-CoV-2 genome Assembler: the ESCA pipeline. Results showed that ESCA can perform high quality genome assembly from IonTor-rent and Illumina raw data, and help the user in easily correct low-coverage regions. Moreover, ESCA includes the possibility to compare assembled genomes of multi sample runs through an easy table format.

Script and manuals are available on GitHub: https://github.com/cesaregruber/ESCA

## Introduction

Whole genome sequence NGS have reached a pivotal role in emerging infectious diseases field, i.e. enhancing development capacity of new diagnostic methods, vaccines and drugs [1,2], and a key role have been recognized to sequence data production and sharing in outbreak response and management [3–5]. In the current COVID-19 epidemic, more than one million of full genome sequences of severe acute respiratory syndrome coronavirus 2 (SARS-CoV-2) have been deposited in publicly accessible data-bases within one year (i.e. GIDAID) [6,7]. SARS-CoV-2 genomes surveillance at global scale is permitting a real-time analysis of outbreak, with a direct impact on the public health response. This contribution comprehends the spread tracing of SARS-CoV-2 over time and space, evidencing emerging variants that may influence pathogenicity, transmission capacity, diagnostic methods, therapeutics, or vaccines [8–11]. Recently divergent SARS-CoV-2 variants are emerging in rapid succession, harboring multiple deletions and amino acid mutations. Some mutations occur in receptor-binding domain of the spike protein and are associated with an increase of ACE2 affinity as well as to hamper recognition by polyclonal human plasma antibodies [12, 13]. The growing contribution of sequence information to public health is driving global investments in sequencing facilities and scientific programs [14,15]. The falling cost of generating genomic NGS data provides new chances for sequencing capacity expansion; however many laboratories have low sequencing capacity and even a lack of data elaboration experties.

While the sequencing runs can be performed without a consolidated experience in infectious diseases field, the virus genomic sequences assembly is often a demanding task. Translating SARS-CoV-2 raw reads data int reliable and informative results is complex, and requires solid bioinformatics knowledge, particularly for low coverage regions that can bring to incorrect variant calling and produce erroneous assembled sequences.

Supervision of sequence assembly to avoid inconsistent or misleading assignment of virus lineage and clade [9, 10], as well as evaluation of low coverage samples to prevent loss of epidemiological information are mandatory.

To this respect, we propose the Easy-to-use SARS-CoV-2 Assembler pipeline (ESCA): a novel reference-based genome assembly pipeline specifically designed for SARS-CoV-2 data analysis. This pipeline was created to support laboratories with low experience in bioinformatics or in SARS-CoV-2 analysis. ESCA can be easily installed and runs in most Linux environments.

## Methods

ESCA pipeline is a reference-based assembly algorithm, written for Linux environments, requires only raw reads as input files, without any other information. Two versions of the software are available: for Illumina pairedend reads in “fastq.gz” file format, and for IonTorrent reads in “ubam” file format.

The software is designed to process several samples in a single run. All reads (paired or unpaired) must be copied into the same working directory, and then launched the program through command line by digiting “StartEasyTorrent” for IonTorrent input or “StartEasyIllumina” for Illumina input. The pipeline than performs automatically all other passages described below. The program processes all input reads, dividing them in different samples using file names as identifiers; Illumina paired-end reads are expected to be divided into two files, with file names that start with the same first 5 letters and that contain “R1” or “R2” to distinguish between forward reads and reverse reads.

Sample preprocessing is performed filtering out all reads with mean Phred quality score lower than 20 and less than 30 nucleotides long. Filtered reads are mapped on SARS-CoV-2 reference genome Wu-han-Hu-1 (GenBank Accession Number NC_045512.2) with bwamem software [15]; all reads that do not map on the reference ge-nome are then discarded.

Genome coverage is then analyzed: read-mapping file is converted in “sorted-bam” and “mpileup” files using samtools software [16] and those data are translated in a detailed coverage table that reports the count of nucleotides observed at each position.

Consensus sequence is then reconstructed based on three parameters:

1. The frequency of nucleotides observed at each position.
2. The nucleotide coverage.
3. The reference genome sequence.

Briefly, sample parameters for consensus sequence reconstruction are designed to call the nucleotide observed with frequency >50% and with a coverage >50 reads; but the minimum coverage is reduced at >10 reads if the most frequent nucleotide observed is identical to the nucleotide observed into the reference genome. For all positions where the parameters described above are not satisfied, ESCA pipeline is designed to call “N” for indicating a low coverage position or an intra-sample nucleo-tide variant.

After whole genome reconstruction of all samples, the consensus sequences are aligned with Wuhan-Hu-1 reference genome, using MAFFT software [17], and a mutation table is generated, reporting nucleotide mutations of all genomes assembled.

To test the efficiency of the EASY, 228 SARS-CoV-2 positive samples were sequenced with Ion Torrent or Illumina platforms using Ion AmpliSeq SARS-CoV-2 Research Panel, following manufacturer’s instructions (ThermoFisher).

For Illumina samples, whole SARS-CoV-2 genome sequence were assembled using both ESCA and DRAGEN RNA Pathogen Detection v.3.5.15 (BaseSpace), with default parameters. For IonTorrent samples, whole SARS-CoV-2 genome sequences were assembled with ESCA and IRMA software [18], using the setting parameters indicated by ThermoFisher. The respectively results were compared aligning the sequences obtained with the two methods with the reference (Wuhan-Hu-1, NCBI Acc.Numb. NC_045512.2) and the corrected sequence submitted to GISAID, using MAFFT [19]. Each discordant position was then evaluated following the classification reported in Fig 1. In particular: TP, true positives (mutation correctly classified as real); FN, false negatives (mutations correctly classified as unreal); FP, false positives (mutations incorrectly classified as real), and TN be true negatives (mutations correctly classified as unreal); T Ncorrected (positions unknown correctly classified as N); T Nerr (positions unknown uncorrectly classified as N).

**Figure 1:**
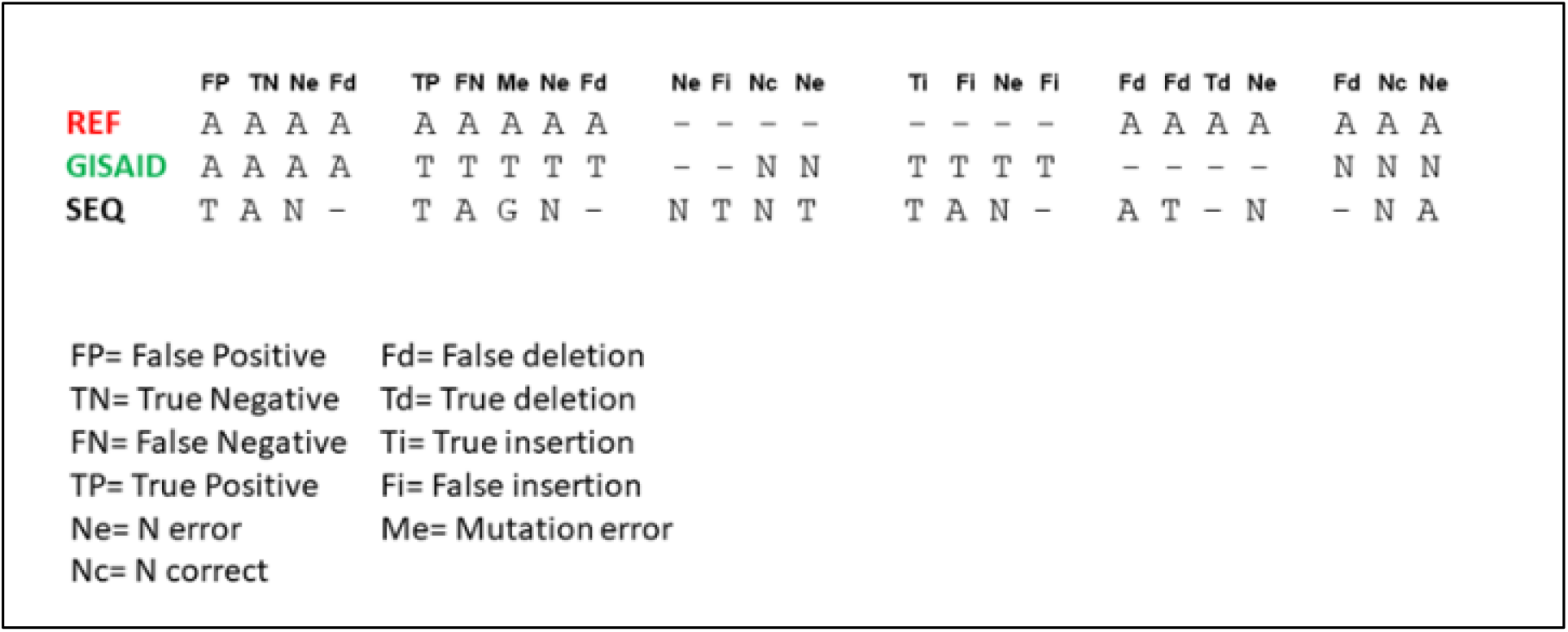
Classification scheme of ge-nome assemblers. Assembled genome sequences (SEQ) were compared with the corresponding submitted sequences (on GISAID), and with reference genome sequence “Wuhan-Hu-1” (REF). Nucleotide threesomes were classified in the eleven categories reported below.

To test the performances with respect to mean coverage, linear regression correlation was carried out between mean coverage and specific measures of accuracy.

## Results and Discussion

In the computational evaluation, ESCA software was compared with the most used assemblers for SARS-CoV-2 genome analysis, on 228 SARS-CoV-2 positive samples. Sequencing was performed on Illumina MiSeq for 65 libreries, obtaining a median of 1.50 x106 paired-end reads per sample (range: 0.02×106 – 4.56×106); and on Ion Gene Studio S5 Sequencer for 163 libraries, obtaining a median of 0.61×106 single-end reads per sample (range: 0.02×106 – 3.02×106). Using the ESCA reconstruction, we calculate the coverage point by point and observed that in illumine sample the point coverage was not uniform, although the mean coverage is quite high in all sample (average 3508X with a range of 70-10733). This could introduce error in genome reconstruction using some software. In this contest Easy could reduce the error in regions with low coverage. In parallel, coverage obtained with IonTorrent was in mean 4966X with a range 94-19917, but a higher uniformity was observed. The comparison of coverage distribution was showed in Fig 2.

**Figure 2:**
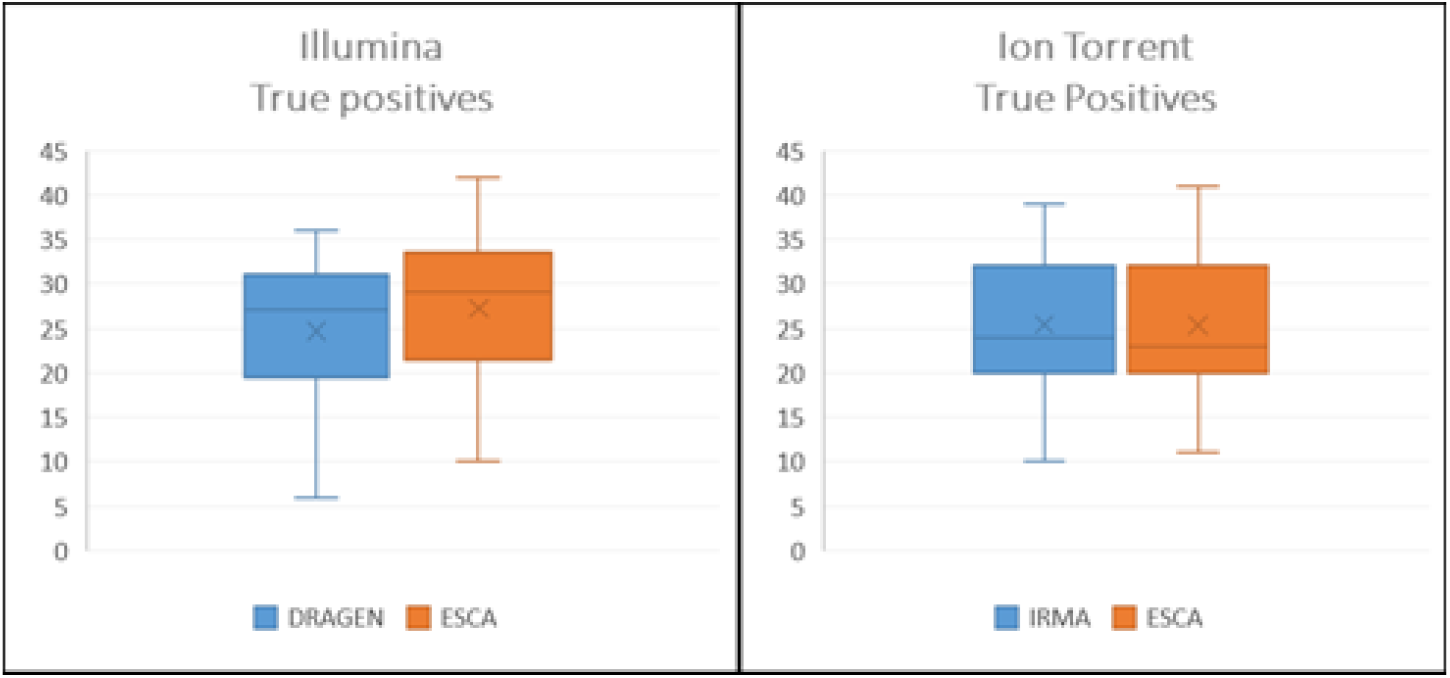
Comparison of True Positive mutations. Assembled genome sequences (SEQ) were compared with the corresponding submitted sequences (on GISAID), and with reference genome sequence “Wuhan-Hu-1” (REF). Nucleotide three-somes were classified in the eleven categories below.

To evaluate the ESCA and Dragen/IRMA results, assembled ge-nomes, reference wuhan-Hu-1 and corrected genome of GISAID (Accession IDs available in Supplementary Table 1) were aligned with MAFFT [19]. At each position along SARS-CoV-2 genome, the 24 available nucleotide combinations were classified in 11 mutation categories (Fig. 1). For all sequences were then evaluated the number of occurrences of mutation categories for each assembly software.

### Illumina data analysis

The comparison of ESCA vs Dragen showed that, as expected, the mean number of mutations in genomes were very low (in mean 28 position) and ESCA could identify correctly 27/28 mutations in mean (Fig 2a). Moreover, no FN positions were identified to ESCA. This is due to the pipeline design that reduce the error introducing N where the coverage was not sufficient. The Dragen genome, instead, showed 25/28 TP and 3 FN positions in mean. The absence of mutation in specific position could be essential to assigned the linage and the presence of false negative could modify the identification of the variants. On the other hand, both ESCA and Dragen not introduce FN, identifying respectively 29308 and 28027 TN positions. These results showing an accuracy of 100.00% in ESCA and 99.99% in Dragen. Moreover, the sensitivity in ESCA vs Dragen was 96.43% vs 89.29% respectively and the specificity in both methods were 100.00%.

### IonTorrent data analysis

Parallel to the previous comparison, ESCA vs IRMA showed that both methods identify 25/26 TP positions in mean (Fig 2b), but not induce FN. However, IRMA introduce some errors, in fact, the FP in IRMA were 20 vs 0 in ESCA. Once again, the introduction of mutations could induce error in lineage assignation, similarly to the absence on real mutations. The accuracy in IRMA was calculated as 99.93% vs 100.00% of ESCA. Moreover, while the sensitivity was identical in the two methods (96.15%) the specificity was 99.93% vs 100.00% in IRMA and ESCA respectively.

### Performance test comparison

To evaluate the performances of each methods, linear regression correlation was carried out with respect to mean coverage (Supplementary Table 2). For IonTorrent single-end sequencing data the significant positive correlation was found between coverage and TN both for IRMA and ESCA (r > 0.15, p < 0.05), while for Illumina pair-end sequencing data such correlation was found only for DRAGEN (r > 0.40, p < 0.05). This difference could be caused by different error rate between two sequencing techniques. These data suggest that all assembly methods are comparable in case of high coverage samples, while ESCA seems to maintain better performance also in low coverage data.

## Conclusion

The importance to rapidly obtain and share high quality whole ge-nomes of SARS-CoV-2 is increasing with the emerging variant strains [14]. For this reason, use of customed ampliconbased sequencing kits can be a rapid and performing method to identify viral variants. However, the lack of bioinformatic skills could be a trouble to handle NGS raw data. Our pipeline ESCA, provides to help laboratories with low bioinformatics capacity, using a single command. Both the methods more common in the analysis of IonTorrent and Illumina data, IRMA and Dragen respectively, shown some error that could induce false identification in variants assignation. On the contrary, SARS-CoV-2 genome obtained by ESCA shows a reduced number of false insertions, false mutations and a higher number of real mutations. Moreover, ESCA make automatically a variant table output file, fundamental to rapidly recognize variants of interest. All these results shown how ESCA could be a useful method to obtain a rapid, complete and correct analysis also with minimal skill in bioinformatics.

## Acknowledgements

We gratefully acknowledge the contributors of genome sequences of the newly emerging coronavirus, i.e. the Originating Laboratories, in sharing sequences and other metadata through the GISAID Initiative, on which this research is based. We also acknowledge Ornella Butera, Francesco Santini e Giulia Bonfiglio for their contribution in sample preparation.

## References

[1] WHO. Genomic sequencing of SARS-CoV-2: a guide to implementation for maximum impact on public health. ISBN 978-92-4-001844-0.

[2] Greaney, A.J., Loes, A.N., Crawford, K.H.D., Starr, T.N., Malone, K.D., Chu, H.Y., Bloom, J.D., Comprehensive mapping of mutations in the SARS-CoV-2 receptor-binding domain that affect recognition by polyclonal human plasma antibodies, Cell Host and Microbe (2021), doi: https://doi.org/10.1016/j.chom.2021.02.003.

[3] Smith GJD, Vijaykrishna D, Bahl J, Lycett SJ, Worobey M, Pybus OG, et al. Origins and evolutionary genomics of the 2009 swine-origin H1N1 influenza A epidemic. Nature. 2009;459:1122–5. doi: 10.1038/nature08182.

[4] Revez J, Espinosa L, Albiger B, Leitmeyer KC, Struelens MJ; ECDC National Microbiology Focal Points and Experts Group. Survey on the Use of Whole-Genome Sequencing for Infectious Diseases Surveillance: Rapid Expansion of European National Capacities, 2015-2016. Front Public Health. 2017 Dec 18;5:347. doi: 10.3389/fpubh.2017.00347. PMID: 29326921; PMCID: PMC5741818.

[5] Nadon C, Van Walle I, Gerner-Smidt P, Campos J, Chinen I, Concepcion-Acevedo J, Gilpin B, Smith AM, Man Kam K, Perez E, Trees E, Kubota K, Takkinen J, Nielsen EM, Carleton H; FWD-NEXT Expert Panel. PulseNet In-ternational: Vision for the implementation of whole genome sequencing (WGS) for global food-borne disease surveillance. Euro Surveill. 2017 Jun 8;22(23):30544. doi: 10.2807/1560-7917.ES.2017.22.23.30544. PMID: 28662764; PMCID: PMC5479977.

[6] Elbe, S.; Buckland-Merrett, G. Data, disease and diplomacy: GISAID’s innovative contribution to global health. Glob. Chall. 2017, 1, 33–46.

[7] NCBI Resource Coordinators. Database resources of the National Center for Biotechnology Information. Nucleic Acids Res. 2018 Jan 4;46(D1):D8–D13. doi: 10.1093/nar/gkx1095. PMID: 29140470; PMCID: PMC5753372.

[8] Dong E, Du H, Gardner L. An interactive web-based dashboard to track COVID-19 in real time. Lancet Infect Dis. 2020 May;20(5):533–534. doi: 10.1016/S1473-3099(20)30120-1. Epub 2020 Feb 19. Erratum in: Lancet In-fect Dis. 2020 Sep;20(9):e215. PMID: 32087114; PMCID: PMC7159018.

[9] Hadfield J, Megill C, Bell SM, Huddleston J, Potter B, Callender C, Sagulenko P, Bedford T, Neher RA. Nextstrain: real-time tracking of pathogen evolution. Bioinformatics. 2018 Dec 1;34(23):4121–4123. doi: 10.1093/bioinformatics/bty407. PMID: 29790939; PMCID: PMC6247931.

[10] Rambaut A, Holmes EC, O’Toole Á, Hill V, McCrone JT, Ruis C, du Plessis L, Pybus OG. A dynamic nomenclature proposal for SARS-CoV-2 lineages to assist genomic epidemiology. Nat Microbiol. 2020 Nov;5(11):1403–1407. doi: 10.1038/s41564-020-0770-5. Epub 2020 Jul 15. PMID: 32669681; PMCID: PMC7610519.

[11] Julia L. Mullen, Ginger Tsueng, Alaa Abdel Latif, Manar Alkuzweny, Marco Cano, Emily Haag, Jerry Zhou, Mark Zeller, Nate Matteson, Kristian G. An-dersen, Chunlei Wu, Andrew I. Su, Karthik Gangavarapu, Laura D. Hughes, and the Center for Viral Systems Biology outbreak.info. Available online: https://outbreak.info/ (2020)

[12] Greaney AJ, Loes AN, Crawford KHD, Starr TN, Malone KD, Chu HY, Bloom JD. Comprehensive mapping of mutations in the SARS-CoV-2 recep-tor-binding domain that affect recognition by polyclonal human plasma anti-bodies. Cell Host Microbe. 2021 Mar 10;29(3):463–476.e6. doi: 10.1016/j.chom.2021.02.003. Epub 2021 Feb 8. PMID: 33592168; PMCID: PMC7869748.

[13] Focosi D, Maggi F. Neutralising antibody escape of SARS-CoV-2 spike pro-tein: Risk assessment for antibody-based Covid-19 therapeutics and vac-cines. Rev Med Virol. 2021 Mar 16. doi: 10.1002/rmv.2231. Epub ahead of print. PMID: 33724631.

[14] WHO. SARS-CoV-2 genomic sequencing for public health goals: Interim guidance. WHO/2019-nCoV/genomic_sequencing/2021.1

[15] Li H, Durbin R. Fast and accurate short read alignment with Burrows-Wheeler transform. Bioinformatics. 2009 Jul 15;25(14):1754–60. doi: 10.1093/bioinformatics/btp324. Epub 2009 May 18. PMID: 19451168; PMCID: PMC2705234.

[16] Li H, Handsaker B, Wysoker A, Fennell T, Ruan J, Homer N, Marth G, Abe-casis G, Durbin R; 1000 Genome Project Data Processing Subgroup. The Se-quence Alignment/Map format and SAMtools. Bioinformatics. 2009 Aug 15;25(16):2078–9. doi: 10.1093/bioinformatics/btp352. Epub 2009 Jun 8. PMID: 19505943; PMCID: PMC2723002.

[17] Katoh K, Misawa K, Kuma K, Miyata T. MAFFT: a novel method for rapid multiple sequence alignment based on fast Fourier transform. Nucleic Acids Res. 2002 Jul 15;30(14):3059–66. doi: 10.1093/nar/gkf436. PMID: 12136088; PMCID: PMC135756.

[18] Shepard SS, Meno S, Bahl J, Wilson MM, Barnes J, Neuhaus E. Viral deep sequencing needs an adaptive approach: IRMA, the iterative refinement meta-assembler. BMC Genomics. 2016 Sep 5;17(1):708. doi: 10.1186/s12864-016-3030-6. Erratum in: BMC Genomics. 2016 Oct 13;17 (1):801. PMID: 27595578; PMCID: PMC5011931.

[19] Katoh K, Misawa K, Kuma K, Miyata T. MAFFT: a novel method for rapid multiple sequence alignment based on fast Fourier transform. Nucleic Acids Res. 2002 Jul 15;30(14):3059–66. doi: 10.1093/nar/gkf436. PMID: 12136088; PMCID: PMC135756.

